# CRISPR/Cas9 screens Reveal Dasatinib Targets of Inhibiting T cell Activation and Proliferation

**DOI:** 10.1101/448662

**Authors:** Mei Wang, Qilong Chen, Hailing Liu, Hao Chen, Binbin Wang, Wubing Zhang, Ailu Mading, Wenjing Li, Shuangqi Li, Hang Liu, ZhongKai Gu, Shuxian Liu, Rui Guo, Wei Li, Tengfei Xiao, Feizhen Wu

## Abstract

Immune response by T cells is essential for a healthy body against cancer, infection, and pathophysiological alteration. The activation and expansion of T cells can be inhibited by dasatinib, a tyrosine inhibitor, thus improving the outcome of diseases, such as autoimmune disease, graft-versus-host disease, and transplant rejection. The underlying mechanism of inhibition by dasatinib is elusive. Here, we designed and synthesized a CRISPR/Cas9 screening library that includes 6,149 genes. Using the library, we performed dasatinib CRISPR/cas9 screening in Jurkat cell, a T lymphocyte cell. We firstly identified survival essential genes for Jurkat cells. Comparing with other CRISPR/Cas9 screenings, we obtained Jurkat cell specific essential genes. By comparing dasatinib treatment to control, we identified a set of dasatinib targets, which includes known targets: CSK, LCK, ZAP70, and previously unknown targets: ZFP36L2, LRPPRC, CFLAR, PD-1, CD45 et al. Visualizing these target genes on T cell receptor signaling pathway, we found several genes could be inhibited by dasatinib. Furthermore, we introduced a framework, 9-square, to classify genes and found a group of genes that are associated with dasatinib resistance, possibly linking the side effects of dasatinib. These data reveal a set of dasatinib targets and demonstrate the molecular potential functions of dasatinib. Identification of dasatinib targets will broaden our understanding to its molecular mechanism, and thus benefits to clinical outcome.

## Introduction

T cells are critical for human against cancer, infection, and autoimmune diseases. Immune responses by T cells are tightly regulated by pathogen-, tumor-antigens, and self-peptides [1]. The activations of T lymphocytes are intimately implicated in several pathophysiological processes of diseases, such as cancer, infection, autoimmune, graft-versus-host disease (GVHD), and transplant rejection. The activations are determined by the major histocompatibility complex (MHC) that binds to antigens derived from pathogens and displays them on the cell surface for recognition by T cells that express appropriate T cell receptor (TCR). The interaction between TCR and MHC along with additional potential signal transduction networks determines the fate of immune response. It is established that 4 families of protein tyrosine kinases: Src family kinases, ZAP-70/Syk, Tec, and Csk, are involved in TCR signaling. Two Src kinases, Lck and Fyn, involving in the initial process of TCR activation, phosphorylate immunoreceptor tyrosine-based activation motifs (ITAMs) in the cytoplasmic domains of CD3- and z-chains within the TCR:CD3 complex. This phosphorylated ITAMs recruit Zap70 that is further activated by Lck-mediated phosphorylation, fully activated Zap70 to form an effective signaling complex, which recruits a number of signaling proteins, including GADS, Grb2, SOS, Vav1, ITK, and PLC-γ1 [2]. The activations of T cells by the TCR signaling initiate a series of complicated biochemical cascades. However, if the activation of the T cell is aberrant, the immune system could be over-response to refrain from protecting and even attacking the body in diseases, such as autoimmune disease, GVHD, and transplant rejection. In such cases, the immune responses are required to be mitigated by inhibiting the activations of T cells with agents.

Dasatinib, a second-generation tyrosine kinase inhibitor (TKI), inhibits T cell activations by targeting multiple kinases, such as Src family. Dasatinib as salvage therapy for sclerotic chronic graft-vs-host disease was reported. Dasatinib is a first-line therapy for chronic myelogenous leukemia (CML) patients [3]. CML is a hematopoietic stem cell disorder frequently caused by chromosomal abnormality, called Philadelphia (Ph) chromosome, a translocation between the abelson (ABL) tyrosine kinase gene at chromosome 9 and the breakpoint cluster region (BCR) at chromosome 22, which forms a fusion oncogene BCR-ABL. Dasatinib is a potent multi-kinase inhibitor that can target and inhibit the BCR-ABL, SRC family (SRC, LCK, HCK, YES, FYN, FGR, BLK, LYN, FRK), receptor tyrosine kinases (c-KIT, PDGFR, DDR1/2, c-FMS, ephrin receptors), and TEC family kinases (TEC and BTK). The Src family kinases are involved in T cell activation, signal transductions, blocking cell duplications, adhesion et al [6]. Dasatinib potently inhibits Src family kinases, diminishing metastatic spread of tumor cells, trigging apoptosis of tumor cells, and inhibiting T cell over-activation [7]. These discoveries suggest that dasatinib can inhibit multiple targets and consequently determine cancer cell fate.

The CRISPR/Cas9 is a gene-editing technology that can cause double-strand breaks (DSBs) by single-guide RNAs (sgRNAs) binding. A sgRNA usually contains 18-20 nucleotides complementary to its target and guides the Cas9 enzyme to a specific DNA location where Cas9 induces a DSB. Repairs of such DSBs by the cell induce mutations in sgRNA-targeted genes, leading to knockouts of these targeted genes[8]. Genome-wide or pooled CRISPR/Cas9 screening is a high-throughput technology to investigate new biological mechanisms, such as drug resistance and cell survival-essential genes. In the screening experiment, sgRNAs are designed, synthesized, and cloned into a lentivirus library, which is subsequently transfected into cells at a low multiplicity of infection to ensure each cell takes up only one sgRNA. Cells are cultured under different experimental settings, such as with or without drug treatment. The sgRNAs that incorporated into the host genome are replicated with the host cell division during passages. By sequencing sgRNAs between initial cells and survived cells using high-throughput sequencing and data analysis, survival essential candidate genes can be identified in comparison between sgRNA in survived cells and initial cells. If one cell carrying a sgRNA is dead or slow-growing, the gene targeted by the sgRNA is possibly essential for the cell survival. Drug-targeting genes can be identified by comparison of CRISPR screenings between with and without drug treatment. Positively selected genes induce drug-treated cells growing faster than control, might be associated with drug resistance, while negatively selected genes induce cell death or growing slowly, might be promising candidates of the drug target. The CRISPR/Cas9 screening has been successfully used for identifying cell-essential, non-essential genes, and drug-targeted genes in genome-scale [9–12].

Despite these progress in finding the regulators for T cell activation and proliferation during the past 20 years, the detailed molecular mechanism underlying TCR signaling for T cell function remains elusive. To address this, we designed and synthesized a gene-pooled CRISPR/Cas9 screening library, which includes 6,149 genes. Using the library, we performed CRISPR/Cas9 screening in human Jurkat cells with dasatinib treatment. By bioinformatics analysis, we identified survival essential genes for Jurkat cells and found the positively selected genes were significantly enriched in the TCR signaling pathway. By comparing with other more than 33 CRISPR/Cas9 screenings, we obtained Jurkat cell specific essential genes. Comparing dasatinib treatment to control, we identified a set of dasatinib targets. Importantly, we introduce a framework, 9-square, to classify screening genes and define a group of genes that are associated with dasatinib resistance and possibly links the side effects of dasatinib. Collectively, our results reveal a group of dasatinib-targets, demonstrating the potential dasatinib mechanism on inhibition of T cell activation and proliferation. Deciphering dasatinib molecular basis is significantly benefit to patients with diseases, such as autoimmune, GVHD, transplant rejection, and CML. These data extend our knowledge regarding the dasatinib molecular mechanism.

## Results

### CRISPR/Cas9 Screening with a custom library

To identify the dasatinib targets with CRISPR/Cas9 screen, we designed and synthesized a custom CRISPR/Cas9 library. Using the library, we performed CRISPR/Cas9 screens with dasatinib treatment in Jurkat cells. Jurkat cells are an immortalized line of human T lymphocyte cells. The library contained 6,149 genes with 63,949 sgRNAs, ~10 sgRNAs per gene. The number of sgRNAs per gene is greater than that in GeCKO (v2) libraries that contains ~6 sgRNAs per gene [11]. Screening results from more sgRNAs per gene are usually more robust than less sgRNAs per gene. Of the 6,149 genes, there are three genomic safe harbors: AAVS1, ROSA26, and CCR5 that are non-essential targeting regions, which are treated as a positive controls for data normalization, because depletion of these genes could not obviously influence cell growth [13]. These 6,149 genes were involved in several important KEGG pathways, including ribosome, cell cycle, pathways in cancer, CML, AML, and et al (Fig 1b). All sgRNAs were synthesized and cloned into a lentivirus library that was then infected into cells at a low multiplicity of infection to ensure each cell takes up only one sgRNA as possible (Fig 1a). The concentration of dasatinib treatment was identified with the half maximal inhibitory concentration (IC50) experiments (Fig 1c). The IC50 of dasatinib was 300 nano-molar in jurkat cells. SgRNAs that were incorporated into the host genome can be replicated with the host cell division. These virally infected cells were selected and split into 3 populations at a starting time point. One population representing the initial sgRNA status (day0) was harvested for sequencing. The remaining 2 populations were respectively treated with dasatinib (treatment) and vehicle (control) and allowed to grow under same experimental conditions for 10 double time (~4 weeks). The screening operations were conducted in parallel to minimize the technical variations. At the end of the screen, genomic DNA from the infected cells was extracted and the sgRNA-encoded regions that viruses had integrated to host genome were sequenced using single-end Illumina platform. 43.67 million reads were generated in the 3 samples in total. The read count of each sgRNA was a proxy for the cell growth characteristics under a knockout of the gene targeted by the sgRNA.

**Figure 1.**
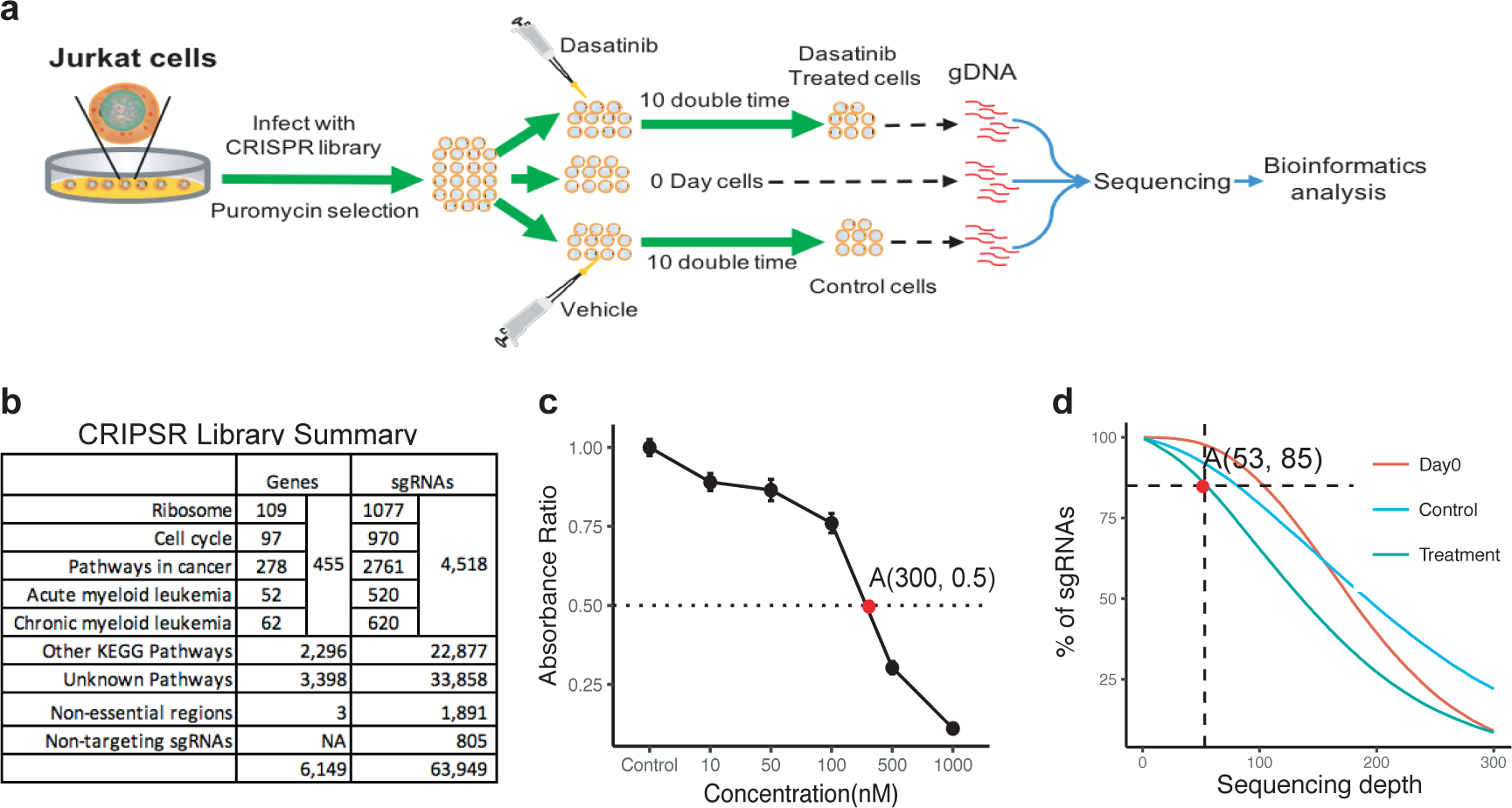
Schema of Dasatinib CRISPR/Cas9 screenings in Jurkat cells. (a) Schema of CRISPR/Cas9 library screening. Jurkat cells were transduced with lentivirus gRNA library, selected with puromysin for 2 weeks, and equally divided into 3 populations: one was treated as base line in a starting time point (Day 0), other 2 populations treated with dasatinib and vehicle as drug treatment and control, respectively. (b) CRISPR/Cas9 library composition. Genes in the library are associated with many essential and cancer related pathways. (c) Half maximal inhibitory concentration (IC50) of dasatinib in jurkat cells. X- and y-axis are dasatinib concentration (nM, nano-molar) and the dasatinib absorbance ratio, respectively. Point A represents the absorbance ratio was 50% when dasatinib concentration was 300nM. (d) sgRNA sequencing depth. In point A, 85% of sgRNAs was sequenced at least 53 converge in day0, control, and treatment samples.

The sequencing data were analyzed using the MAGeCK (v0.5.7) package[14] that uses negative binomial distribution to calculate beta score, which represents CRISPR screening performance on each genes. The beta score is similar to the log2-transferred fold-change in gene differential expression analysis. By comparing dasatinib-treated and vehicle-treated sample to day0 sample, we obtained treatment and control beta score (CBS), respectively. Dasatinib-targeting genes were identified on the basis of comparison of treatment beta scores (TBS) to CBS. Mapping these sequencing reads onto the CRISPR library, we found that at least 85% sgRNAs were sequenced with more than or equal 53 reads in the 3 samples (Fig 1d). 79.4% reads in day0 sample, 81.9% in control, and 81.6% in treatment were mappable (SFig 1a). The mapping rate indicates the successes in construction and sequencing of the CRISPR libraries. Principle component analysis (PCA) and cumulative distribution function (CDF) shown that the 3 samples can be separated from each other and with similar frequency tendency, respectively (SFig 1b, c). Collectively, these data suggest that the 3 samples qualify for downstream analysis with the success in the CRISPR library design, synthesis, screening, sequencing, and bioinformatic analysis.

### Identification of Jurkat-specific essential genes

We firstly identified essential genes for Jurkat cell survival. Identification of essential genes can enable us to understand T cell proliferation. By comparing the control sample to day0, we calculated CBS. Using 1-quantile standard deviation (QSD) as cutoff, we identified 238 positively selected genes (PSGs) with beta scores greater than the cutoff, indicating these gene could suppress T-cell proliferation (Fig 2a and Supplementary table 1). We also identified 1,714 negatively selected genes (NSGs) with beta scores less than the cutoff (Fig 2a and Supplementary table 1), indicating these genes are required for T-cell proliferation. As expected, these NSGs were enriched in ribosome, spliceosome, RNA transport, and cell cycle pathways (SFig2 d, e), indicating these pathways are essential for T-cell proliferation, which suggests the Jurkat cells share common essential genes with other cells. Interestingly, these PSGs were enriched in TCR signaling pathway, insulin resistance, and viral carcinogenesis (SFig2 c). Visualizing of control beta scores in TCR signaling pathway, we found that genes, including CTLA4, CD45, LCK, ZAP70, and et al, were negatively selected, suggesting these genes are essential for T-cell proliferation, while genes, such as NCK, PAK, NFAT, AP1, and et al, were positively selected, suggesting these genes are required to be suppressed for TCR signaling pathway activation. These data suggest these selected genes fine-tune T-cell activation and proliferation through the TCR signaling pathway (SFig2 j). Collectively, we identified positively and negatively selected genes for T-cells and demonstrated these genes are involved in T-cell proliferation.

**Figure 2.**
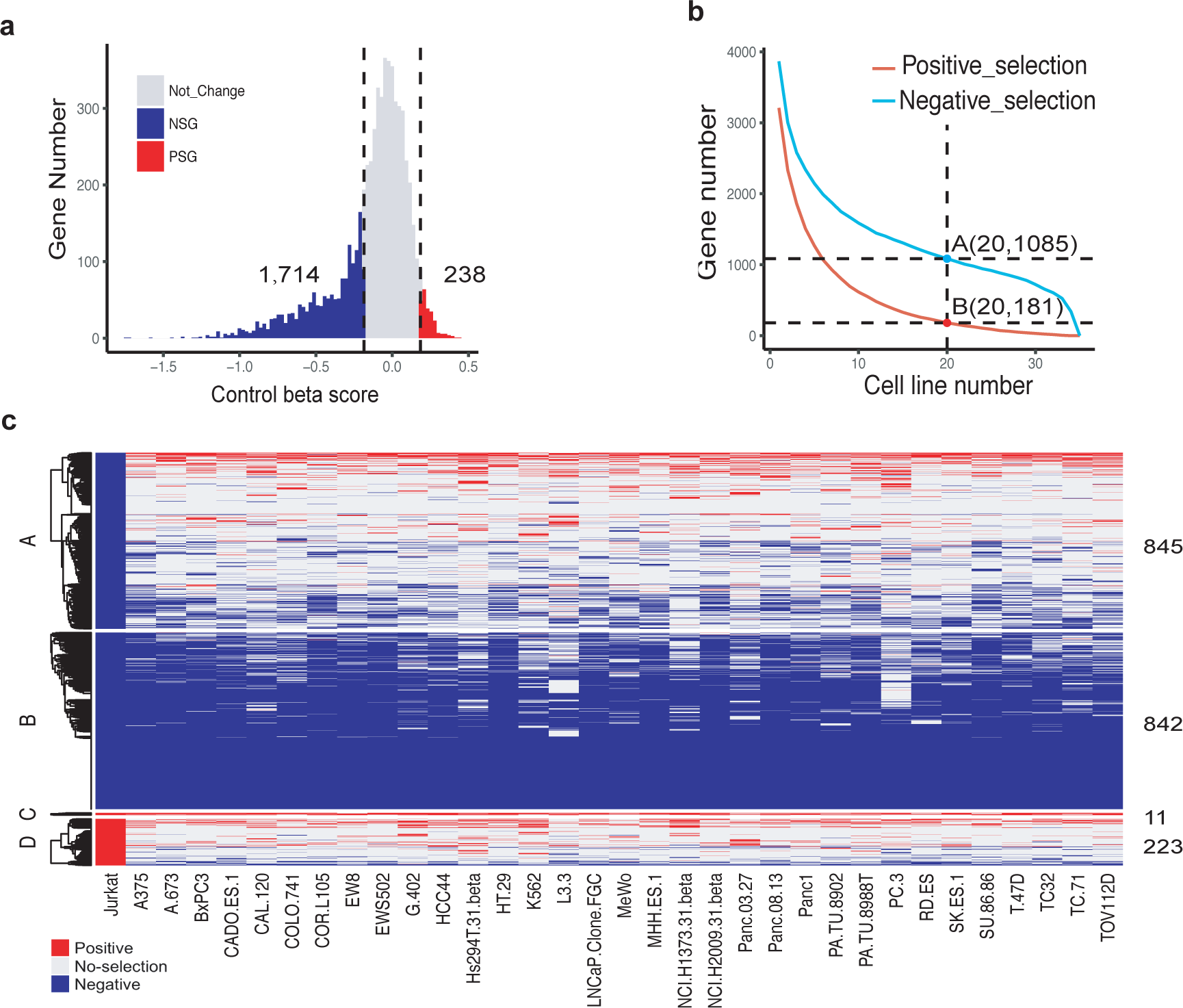
Identification of essential genes in Jurkat cells. (a) A distribution of essential genes of Jurkat cells. 1,714 and 238 are total NSG number and PSG number, respectively. Vertical dotlines are cutoff of +/- 1 QSD. (b) Common positive and negative selected gene numbers across 34 cell lines, including Jurkat. X- and y-axis are the cell line number and common selected-gene number, respectively. Point A represents 1,085 genes were negatively selected across at least 20 cell lines, and point B 181 positively selected across at least 20 cell lines. The 1,085 and 181 genes were defined as common negatively and positively selected genes across 34 cell lines, respectively. (c) A clustering heatmap showing Jurkat positively/negatively selected genes across 34 cell lines. Group A and D were 845 negatively and 223 positively Jurkat-specific selected genes, respectively. Group B and C were 842 negatively and 11 positively commonly selected genes across at least 20 cell lines, respectively. Red and blue are positive and negative selection in the heatmap, respectively. NSG: negatively selected gene; PSG: positively selected gene; QSD: quantile standard deviation.

We next identified genes that are essential for Jurkat cells specifically, which can extend our understanding to the Jurkat cells characteristics. We analyzed CRISPR/Cas9 screening data from a prior study that carried out on 33 cell lines [15] by MAGeCK package. These cell lines represent a wide diversity of cells to a certain extent. We selected 6,089 common screening genes across our Jurkat cells and the 33 cell lines. Among the 33 cell lines, we identified 1,085 NSGs and 181 PSGs across at least more than 20 cell lines at a cutoff of 1 QSD (Fig 2b), suggesting that these genes were selected from the majority of cell lines. Thus, we defined these genes as common positively and common negatively selected genes. Excluding these common positive and negative selection genes from the PSGs and NSGs of Jurkat cells, we obtained 845 Jurkat-specific NSGs (group A) and 223 Jurkat-specific PSGs (group D), as well as 842 common negatively (group B) and 11 positively (group C) selected genes across these cell lines (Fig 2c, SFig 2a, and Supplementary table 2). These 845 NSGs were specifically essential for Jurkat cell survival, reflecting the characteristics of Jurkat cells. We also analyzed the gene expression profile of Jurkat cells using public data in GEO database (GSM2068619 and GSM2068620). Gene expression profile analysis shown that the gene expression abundance in group A and B from Jurkat cells was greater than that in group C, D, and non-selected genes (SFig 2b), and group B greater than group A, which suggests the Jurkat-specific and common negatively selected genes are required high expression relative to other genes for maintaining the cell growth. Performing GO term and KEGG pathway enrichment analysis, we found that these negative Jurkat-specific selected genes were pertaining to such pathways as fanconi anemia pathway, alcoholism, and et al (SFig 2f, g). Visualizing control beta scores onto fanconi anemia pathway, we found almost all screened genes, excepting PMS2, were negative selected, indicating these Jurkat-specific negatively selected genes in fanconi pathway are essential for the cell survival. Taken together, these data suggest that Jurkat cell survival is associated with a group of essential genes, in which the Jurkat-specific selected genes are involved in TCR signaling that possibly links the Jurkat identity.

### Identification of dasatinib-targeting genes

To identify dasatinib-targeting genes, we identified treatment beta scores (TBS) by comparing dasatinib-treated library to day0, and then comparing TBS to CBS. We identified a list of beta score differences between TBS and CBS, which reflects dasatinib influence in the screenings. Similarly, using 1-QSD as a cutoff, we identified 1,053 positively and 902 negatively selected genes (Fig 3a, SFig 3a, and supplementary table 2). By ranking the delta beta scores (treat-control) from high to low, we found that the top 5 genes: CSK (C-Terminal Src Kinase), LCK (LCK proto-oncogene, Src family tyrosine kinase), ZFP36L2 (zinc finger protein 36, C3H type-like 2), LRPPRC (leucine rich pentatricopeptide repeat containing), and CFLAR (CASP8 and FADD like apoptosis regulator), were positively selected. This suggests that the genes more essential in control cells than in dasatinib-treated cells (Fig 3b), indicating that these genes as cell fitness genes whose functions were inhibited by dasatinib treatment. Thus, these genes are potential targets of dasatinib. CSK and LCK as tyrosine kinases, were previously known to be dasatinib targets[16]. By contrast, the bottom 5 genes: SIK3 (SIK Family Kinase 3), CDK6 (cyclin dependent kinase 6), BCL2L1 (BCL2 like 1), KMT2C (lysine methyltransferase 2C), and TONSL (tonsoku like, DNA repair protein), were negatively selected. This suggests that these genes were more essential in dasatinib-treated cells than in control cells (Fig 3b), indicating that the loss of these genes sensitized the cells to dasatinib. Therefore, these genes when inhibited would synergistically enhance dasatinib treatment for T cells. Examining the sgRNA changes between the 3 samples, we found that sgRNAs of LCK, CSK, and CFLAR in dasatinib-treated sample were higher than in control sample, supporting positive selection by dasatinib, while SIK3 and CDK6 in treatment were lower than in control, supporting negative selection by dasatinib (Fig 3c). We performed KEGG pathway enrichment analysis for the bottom 400 negatively selected genes and found these genes were significantly enriched in chronic myeloid leukemia (CML), acute myeloid leukemia (AML), and TGF-beta signaling pathways, suggesting that dasatinib treatment is implicated in the alterations on these pathways (SFig 3b), thus potentially improving the pathology of CML and AML. Performing GSEA analysis, we found many genes in CML and AML pathways were negatively selected (SFig 3d, e, g, h), indicating that CML and AML pathways were suppressed when cells were treated with dasatinib, which supports the therapy of dasatinib in CML and AML patients. More importantly, we found 13 positively and 13 negatively selected genes in TCR signaling pathway at cutoff of 1-QSD (Fig 3d). The positive selected genes include LCK, ZAP70, CD45, GSK3B, CBLB, PD-1, ITK et al, suggesting that the functions of these genes can be inhibited by dasatinib, while IFNG, HRAS, RELA, RAF1, AKT1 et al were negatively selected, indicating these genes could synergistically enhance dasatinib treatment function (SFig 3c). LCK and ZAP70 have previously found to be targets of dasatinib. PD-1 is a checkpoint playing a critical role in regulating TCR signaling pathway. ITK encodes an intracellular tyrosine kinase expressed in T-cells, regulates the development, function and differentiation of T-cells, thus ITK targeted by dasatinib can inhibit T cell activation and proliferation.

**Figure 3.**
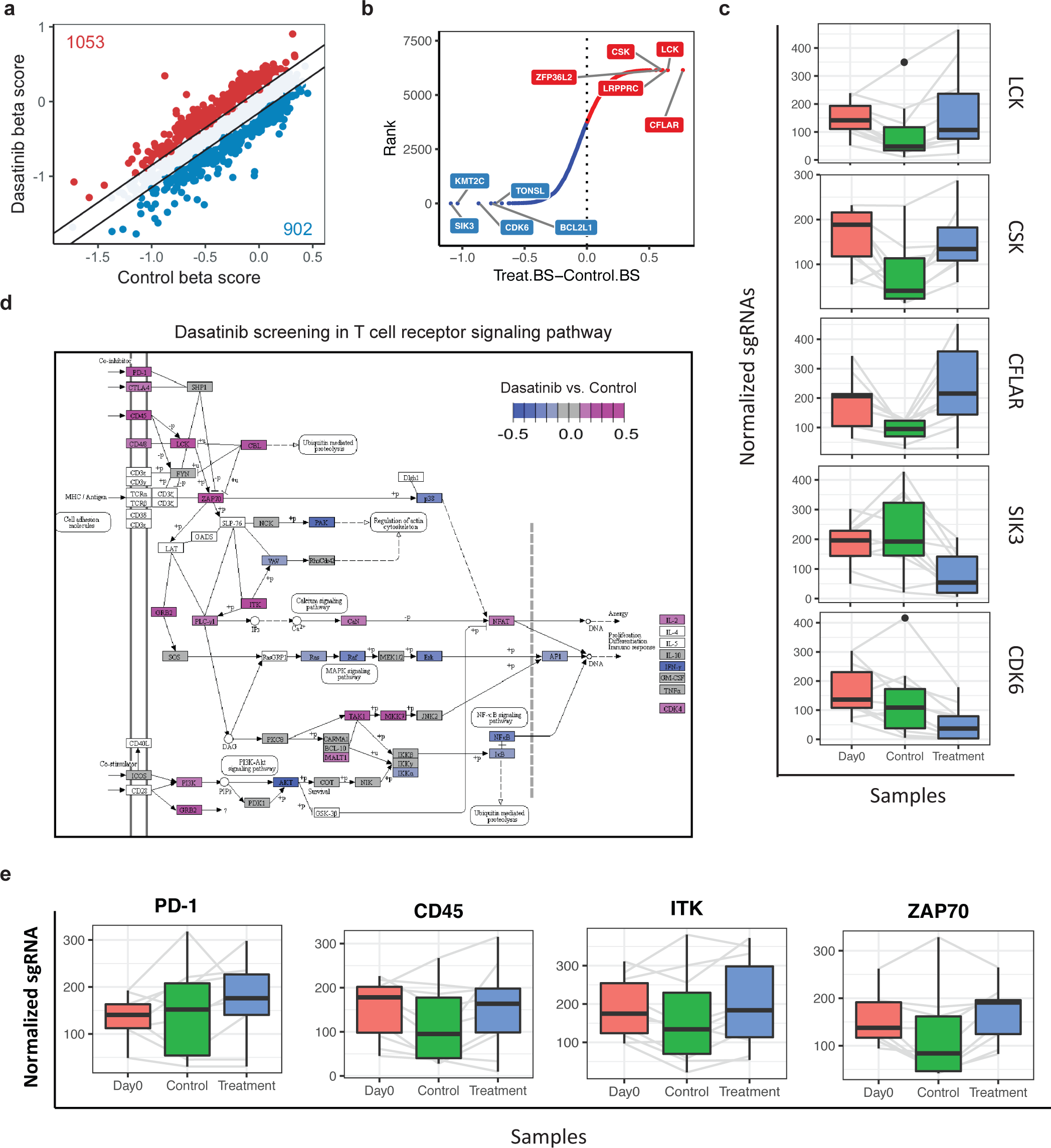
Dasatinib CRISPR screening. (a) A scatterplot shows the distribution of control and dasatinib beta score. (b) Rank of value of treatment minus control beta score. Treat.BS and Control.BS are dasatinib and control beta score. (c) Boxplots show normalized sgRNA count distributions in representive genes: LCK, CSK, CFLAR, SIK3, and CDK6. The normalization is based on the amount of sequencing data. (d) visualization of beta score change of dasatinib/control beta score in T cell receptor signaling pathway. The pathway was adapted from chronic myeloid leukemia KEGG pathway. (e) Boxplots show normalized sgRNA counts in TCR pathway genes: PD-1, CD45, ITK, and ZAP70.

Next, we ask if there were interactions between these identified dasatinib-targets. Based on the annotation of protein-protein interactions (PPI) from GeneMANIA[17], we performed PPI analysis for these dasatinib targets. we highlighted a sub-network that includes ranked top 5 (CSK, LCK, ZFP36L2, LRPPRC, and CFLAR), bottom 5 genes (SIK3, CDK6, BCL2L1, KMT2C, and TONSL), and genes that were interacted with the 10 genes (SFig 3f), suggesting that these dasatinib targets are possibly interacted to mediate dasatinib functions. We, based on these data, concluded that the dasatinib targets the tyrosine kinases and TCR related genes to mediate gene functions, and consequently suppress T cell activation and proliferation.

### Dasatinib resistance and response gene analysis

To further explore dasatinib molecular function, we assigned all genes into 9 squares using 1-QSD as a cutoff for CBS and TBS (Fig 4a). The delta score (TBS-CBS) reflects dasatinib treatment. Genes selected by CBS were associated with cell growth, while genes selected in TBS were associated with both dasatinib treatment and cell growth. We highlighted 4 groups: A, B, C and D (Supplementary Table 3), which were differentially selected between control and treatment screenings. Group A and C were positively and negatively selected in TBS, respectively, but not selected in CBS, suggesting these genes were associated with the drug treatment, but not associated with cell growth. Conversely, group B and D were positively and negatively selection in CBS, respectively, but not selection in TBS, suggesting these genes were associated with cell survival, but not associated with dasatinib treatment. In group A, dasatinib treatment induced cell growing faster than control, indicating genes in group A may be associated with dasatinib resistance, while in group C, cell grew lower than control, indicating may be associated with dasatinib response. Ranking the beta score difference, the top 5 genes in group A are NUDCD1, CST4, DLX5, SIGMAR1, and NLRP7, and bottom 5 genes in group C are SIK3, KMT2C, RPL41, SLC25A22, and PRKDC. We performed GO biological process and KEGG pathway enrichment analysis for group A and C (Fig 4b, c). The top 10 enriched KEGG pathways includes Hedgehog signaling pathway, long-term potentiation, and cholesterol metabolism pathway, and top 10 GO biological processes includes regulation of cell fate commitment, which are consistent with previous studies [18, 19]. Similarly, group B were less positive selection after dasatinib treatment than control, suggesting these genes may be synergy effect with dasatinib, while group D were less essential after dasatinib treatment than control, indicating these genes may be dasatinib targets. We performed enrichment analysis for group B and D as well, indicating that these genes were pertaining into pathways, including systemic lupus erythematosus, viral carcinogenesis, Alcoholism, Th1 and Th2 cell differentiation, et al. Clinical treatment found that CML with long-term taking dasatinib could suffer from side-effects, such as systemic lupus erythematosus, viral infection, increase disease risk when drink alcohol, and immune inhibition of Th1 and Th2. These results provide links to the potential mechanism of the drug side effects. Taken together, Group A and C maybe associated with the dasatinib resistance and response, and group B and D associated with synergy and inhibition effects of dasatinib.

**Figure 4.**
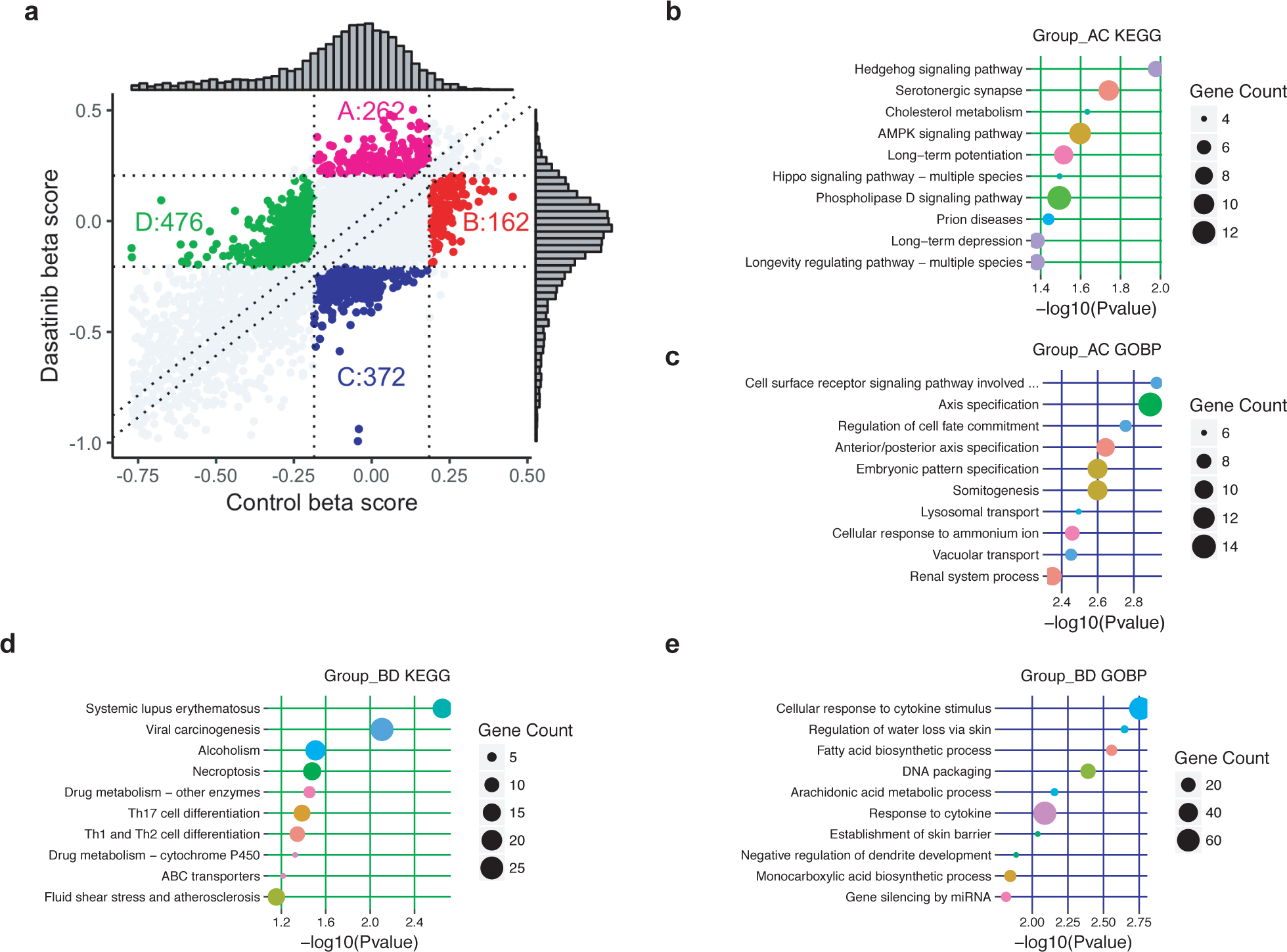
gene classification with control and dasatinib beta score and KEGG and GO enrichment analysis. (a) gene classifications with control and dasatinib beta score. Vertical dot lines are control cutoff on +/- 1 QSD. Horizontal dot lines are dasatinib cutoff on +/- 1 QSD. According to these vertical and horizontal dot lines, all genes are assigned into 9 groups. Diagonal dot lines are lines of dasatinib/control cutoff +/- 0.05 for separating group A and B as well as D and C. (b & c) KEGG pathway (b) and GO biological process enrichment analysis (c) for genes in group A and C. (d & f) KEGG pathway (d) and GO biological process enrichment analysis (f) for genes in group B and D. (e) Values of dasatinib minus control beta score on hedgehog signaling pathway. (g & h) Group A and C gene beta score difference visualization in AMPK signaling pathway (g) and Long-term potentiation (h). (i-k) Group B and D gene beta score difference visualization in system lupus erythematosus (i), viral carcinogenesis (j), and alcoholism (k) KEGG pathways

## Discussion

Our goal is to identify dasatinib targets of inhibiting T-cell activation and proliferation. To achieve these goals, we designed and synthesized a CRISPR/Cas9 screen library, which contains 6,149 genes, covering key cancer-related pathways. SgRNAs in the library were lentivirally transduced into Jurkat cells. 238 and 1,714 genes were respectively positively and negatively selected by comparison to the starting plasmid library after 4-week unperturbed growth. Comparing with 33 published screening datasets, we identified a set of genes that are essential for Jurkat specifically. By comparing dasatinib-treatment to control beta score, we identified a group of potential targets of dasatinib, which are closely associated with TCR, CML and AML pathways. These dasatinib targets included previously known targets: LCK, CSK, and unknown targets: ZFP36L2, LRPPRC, CFLAR et al. Assigning genes into 9-square, we highlighted 4 group genes that are possibly associated with dasatinib resistance, response, and side effects. These findings extend our knowledge regarding T-cell activation, proliferation, and dasatinib molecular function.

T cell activation and proliferation are critical for adaptive immune. Dasatinib can inhibit tyrosine kinase and T cell activation. Although the underlie mechanisms of T cell activation, proliferation, and dasatinib remain to be investigated, several inferences can be drawn from the existing data.

First, we used CRISPR screens to identify 238 positive and 1,714 negative selection genes that are involved in Jurkat cell survival. Jurkat cells are an immortalized line system of human T lymphocyte cells that are used to study acute T cell leukemia, T cell signaling, and the expression of various chemokine receptors susceptible to viral entry, particularly HIV[20]. Jurkat cells can be used to determine the mechanism of differential susceptibility of cancers to drugs. These negatively selected essential genes were enriched in ribosome, spliceosome, and RNA transport pathways, suggesting these genes are strongly associated with T cell survival. These positively selected genes are associated with TCR signaling pathways, indicating that these genes may be associated with the immunomodulation. The positively and negatively selected genes are involved in T cell vulnerabilities. These data facilitate the understanding of T cell activation and proliferation.

Second, to identify Jurkat-specific essential genes, we compared the identified gene set with that from other 33 cell lines. We obtained 845 negatively and 223 positively selected Jurkat-specific genes. The 845 genes are pertaining to Fanconi anemia pathway, Alcoholism, Cysteine and methionine metabolism, and systemic lupus erythematosus pathways, while the 223 associated with B cell and TCR signaling pathway, indicating these pathways may be associated with the identity of Jurkat. Token gather, the Jurkat-specific selected genes will benefit to the community for understanding the T cell characteristics.

Third, as a second-generation TKI inhibitor, the dasatinib is an effective therapy to CML patients carrying a fusion of BCL-ABL. However, the potential mechanism of dasatinib is elusive, requiring further investigations. By comparison of dasatinib-treated screen to control screen, we identified a group of targets that are potentially involved in dasatinib therapy, including CSK, LCK, BCL, and ABL. These genes are targeted and inhibited by dasatinib[7, 21]. These dasatinib-targeted genes were associated with CML, AML, and TCR signaling pathways. Visualizing these targeting genes on KEGG pathways enables intuitive displaying genes in the context of regulation networks.

Fourth, assigning all genes into 9-square with a cutoff of 1-QSD, we identified 4 group genes. Genes positively and negatively selected in dasatinib treatment but not in control may be associated dasatinib resistance and response. These genes are associated with systemic lupus erythematosus, viral carcinogenesis, alcoholism, Th17 cell differentiation, Th1 and Th2 cell differentiation pathways. Clinical evidences shown that patients with long term treatment of dasatinib may suffer severe side effects, including system lupus erythematosus, alcoholism, and immune suppression[22]. These data suggest that these genes may be associated with dasatinib side effects.

Finally, while we believe that these identified genes are important for T cell survival and dasatinib treatment, some issues remain to be addressed. We used Jurkat cells as model, which is different from Ph+ CML cells. The screen library covers 6149 genes only. it is possibly missing some important genes. Although these limitations are existing in the data, these data are largely extending our understanding to dasatinib mechanism, which will facilitate an improvement on dasatinib treatment in clinic. Overall, although our study has certain limitations, because of the cell line in the experiments, requiring additional validations, we nonetheless identified a group of T cell essential genes and found that dasatinib-treatment is closely associated with CML and AML pathways. These findings refine our understanding to T cell proliferation, and dasatinib mechanism, which will facilitate the advantage in the clinic outcome.

## Conclusions

Immune response by T cell activation and expansion is of importance to the body against cancer-antigens, pathogen-, antigens, and self-peptides. if over activation or proliferation of T cell, the immune system could fail to protect and even attack the body with diseases, such as autoimmune disease and GVHD. We have presented CRISPR screens with dasatinib-treatment in T cells. We identified a set of essential genes for T cell survival, which will contribute to the field with extending the understanding of T cell survival mechanism. With dasatinib treatment, we identify dasatinib targets and demonstrating the potential reason of dasatinib inhibition on T cell activation. Collectively, these results and the demonstrations of CRISPR screen have set the stage for further understanding the T cell survival and the mechanism of inhibition by dasatinib on T cell activation and proliferation.

## Acknowledgments

This work was supported by China Scholarship Council (201506105022), The national natural science foundation of china (81100589, 81272295), the national High-tech R&D program, 863 Program (2015AA020108). Xiaole Shirley Liu at Dana-Farber cancer Institute of Harvard University is acknowledged for her expert technical assistance.

## Materials and Methods

### 1. Design and synthesis of CRISPR/Cas9 screening library

To design a smaller scale CRISPR/Cas9 knockout screen library focusing on cancer-related genes, we selected 6000 genes based on reported relevancies with cancers using multiple literature, including Cosmic[23] and Oncopanel [24]. For each gene, we designed ten 19-nucleotide sgRNA against its coding region with optimized cutting efficiency and minimized off-target potentials. We used sequence features of the spacers to calculate the efficiency score for each sgRNA using our predictive model [25] to improve cutting efficiency and utilized Bowtie to map all candidate sgRNAs to hg38 reference genome and chose those with least alignment to reduce potential off-targets. We selected the top 10 best sgRNAs for each gene based on the considerations above. The library also contains both positive controls and two types of negative controls: non-targeting controls and non-essential regions-targeting sgRNAs.

(1) Positive controls: we included 1466 sgRNAs targeting 147 positive control genes, which are significantly negatively selected in multiple screen conditions.

(2) Non-targeting negative controls: 795 sgRNAs with sequences not found in human genome (hg38).

(3) non-essential regions-targeting negative controls: 1891 sgRNAs targeting AAVS1, ROSA26, and CCR5, which have been reported as safe-harbor regions where knock-in leads to few detectable phenotypic and genotypic changes.

| Targets | Cancer – related | Positive controls | Non-targeting | Non-essential targeting regions | Total |
| --- | --- | --- | --- | --- | --- |
| Number of genes | 6000 | 146 | - | 3 | 6149 |
| Number of sgRNAs | 59797 | 147 | 795 | 1891 | 63949 |

### 2. Cell culture and CRISPR screening

#### (1) Cell culture

Jurkat cells were cultured in RPMI1640 (Invitrogen) supplemented with 10% fetal bovine serum (Hyclone) and penicillin/streptomycin and maintained at 37°C in a humidified atmosphere at 5% CO2. The cells were suspension lymphoblasts. When cells were 6-8×105cells/ml, split them 1:4 with fresh media (T75=50ml).

#### (2) Dasatinib treatment

Dasatinib was purchased from Selleck, USA. Dasatinib was prepared as 10mM stock solutions in dimethyl sulfoxide, stored in aliquots at −20 °C until further use. Jurkat cells (5×10^5^ cells/ml) were treated with or without different concentrations of dasatinib about 3 days (0.10, 50, 100, 500, 1000nM). Proliferation was measured after 3 days of culture by CCK8 assay (Dojindo Molecular Technologies, USA).

#### (3) CRISPR screening

1.2×10^8^ Jurkat cells (5×10^5^ cells/ml) were transfected with the customized pooled lentiviral sgRNA library at multiplicity of infection of 0.3. To select for sgRNA-expressing cells, transfected cells were treated with 0.5 µg/mL puromycin, commencing 48h after transduction. Puromycin was withdrawn at day 3, and cells were divided into three groups (Day 0 group, vehicle group, and dasatinib treatment group). The cells in vehicle group were cultured without dasatinib and dasatinib treatment group were cultured with 100nM dasatinib for about 1 month.

#### (4) Genomic DNA sequence

Genomic DNA (gDNA) of three groups was extracted using a Blood & Cell Culture Midi kit (Qiagen). SgRNA-coding regions integrated into the chromosomes were PCR-amplified with Q5 High-Fidelity DNA Polymerase (Biolabs). To keep the fold representation at 300 for each of the sgRNA, 200µg of gDNA per each sample was used for the PCR. SgRNA sequences were amplified by two rounds of PCR. For the first PCR, perform 25-30 separate 100 µl reaction with 6-8 µg gDNA in each reaction and then combined the resulting amplicons. The forward primer of the first PCR is AATGGACTATCATATGCTTACCGTAACTTGAAAGTATTTCG and the reverse primer of the first PCR is TCTACTATTCTTTCCCCTGCACTGTACCTGTGGGCGATGTGCGCTCTG. To attach Illumina adaptors and to barcode samples, the second PCR was performed in a 100 µl reaction volume using 5 µl of the product from the first PCR. Forward primer of the second PCR is AATGATACGGCGACCACCGAGATCTACACTCTTTCCCTACACGACGCTCTTCCGATCTTCTTGTGGAAAGGACGAAACACCG and reverse primer of the second PCR is CAAGCAGAAGA CGGCATACGAGATGTGACTGGAGTTCAGACGTGTGCTCTTCCGATCT (NNNNNNNN, 8 bp-barcode) TACTATT CTTTCCCCTGCACTGTACC. PCR bands were gel purified using Qiagen gel purification kit. The resulting libraries were sequenced with pair-end 150bp reads on an Illumina Hiseq 2500.

### 3. CRISPR screening data analysis

The CRISPR/Cas9 screening data were analyzed using the MAGeCK and MAGeCK-VISPR algorithms [14] [26]. We use the MAGeCK-VISPR algorithm to compare the gene selections across different conditions. MAGeCK-VISPR uses a metric, “β score”, to measure gene selections. The definition of β score is similar to the term of ‘log fold change’ in differential expression analysis, and β>0 (or <0) means the corresponding gene is positively (or negatively) selected, respectively. MAGeCK-VISPR models the gRNA read counts as a negative binormal variable, which mean value is determined by the sequencing depth of the sample, the efficiency of the gRNA, and a linear combination of β scores of the genes. MAGeCK-VISPR builds a maximum likelihood (MLE) model to model all gRNA read counts of all samples, and iteratively estimate the gRNA efficiency and gene β scores using the Expectation-Maximization (EM) algorithm. A detailed description of the MAGeCK-MLE algorithm can be found in the original study[14].

We identified Jurkat essential genes by control sample compared with starting time sample, using MAGeCK, and dasatinib target genes by treatment β scores compared with control β scores. The MAGeCK algorithm works as follows. It first collects read counts of all gRNAs in all conditions from fastq files, and then normalizes the read counts of control and treatment conditions using median normalization. After that, MAGeCK builds a linear model to estimate the variance of gRNA read counts, evaluate the gRNA abundance changes between control and treatment conditions, and assigns a p-value using the Negative Binomial (NB) model. Finally, the selection of genes is evaluated from the rankings of gRNAs (by their p-values) using the α-RRA (α-Robust Rank Aggregation) algorithm. For each gene, α-RRA evaluates the rankings of all its gRNAs, and assigns a lower score (RRA score) if the distribution is more skewed compared with uniform distribution. The statistical significance of the RRA score is evaluated by permutation, and the Benjamini-Hochberg method is used for multiple comparison adjustments. To increase the statistical power, genes that have fewer than 4 gRNAs, or genes that have fewer than 2 significant gRNAs are excluded from the comparison. A detailed description of the MAGeCK algorithm can be found in the original study [14, 26].

Gene enrichment was performed with clusterProfiler package, which allowed to analyze enrichment of GO biological process terms and KEGG pathways. Pathview package was used to visualizing β scores or β score difference on the KEGG pathways. R-scripts were made to visualize enrichment analysis.

